# Coupling of conformational switches in calcium sensor unraveled with local Markov models and transfer entropy

**DOI:** 10.1101/2020.02.25.964353

**Authors:** Tim Hempel, Nuria Plattner, Frank Noé

## Abstract

Proteins often have multiple switching domains that are coupled to each other and to binding of ligands in order to realize signaling functions. Here we investigate the C2A domain of Synaptotagmin-1 (Syt-1), a calcium sensor in the neurotransmitter release machinery and a model system for the large family of C2 membrane binding domains. We combine extensive molecular dynamics (MD) simulations and Markov modeling in order to model conformational switching domains, their states and their dependence on bound calcium ions. Using transfer entropy, we characterize how the switching domains are coupled via directed or allosteric mechanisms and give rise to the calcium sensing function of the protein. Our switching mechanism contributes to the understanding of the neurotransmitter release machinery and the proposed methodological approach serves as a template to analyze conformational switching domains and study their coupling in macromolecular machines in general.

## 1 Introduction

Molecular modeling comes with challenges. Even though the advent of graphical processing units (GPUs) has delivered vast amounts of data from increasingly large molecular systems, modelers still have to cope with two fundamental problems: The sampling problem and the curse of dimensionality. In particular, a limiting factor of molecular modeling is the (possibly exponential) growth of the number of metastable states with system size. Sampling all metastable states reversibly can thus be unfeasible, especially if the described processes are extremely slow. Recently, dynamical graphical models have been proposed to overcome this problem by treating the evolution of configurations of many small molecular switches, e.g., dihedral rotamers, in an Ising-model like fashion.^1^

Here we develop a complementary approach that, instead of working with very small spin-like molecular switches that can be readily identified from the structure, we identify conformational switching domains and model their kinetics with local Markov state models (MSMs). To this end, we partition the protein into subsystems that are modeled separately as conformational switching domains or, short, conformational switches. We therefore do not need to parametrize an MSM of the full structure and thus require less extensive molecular dynamics (MD) sampling. This decomposition approach is supported by experimental evidence for “independent dynamic [protein] segments”^2^ that have undergone evolutionary development in an independent fashion.^2^ Further, we quantify the coupling between the local conformational switches by using transfer entropy in the time series of Markov states. Our approach is summarized in Fig. 1.

**Figure 1:**
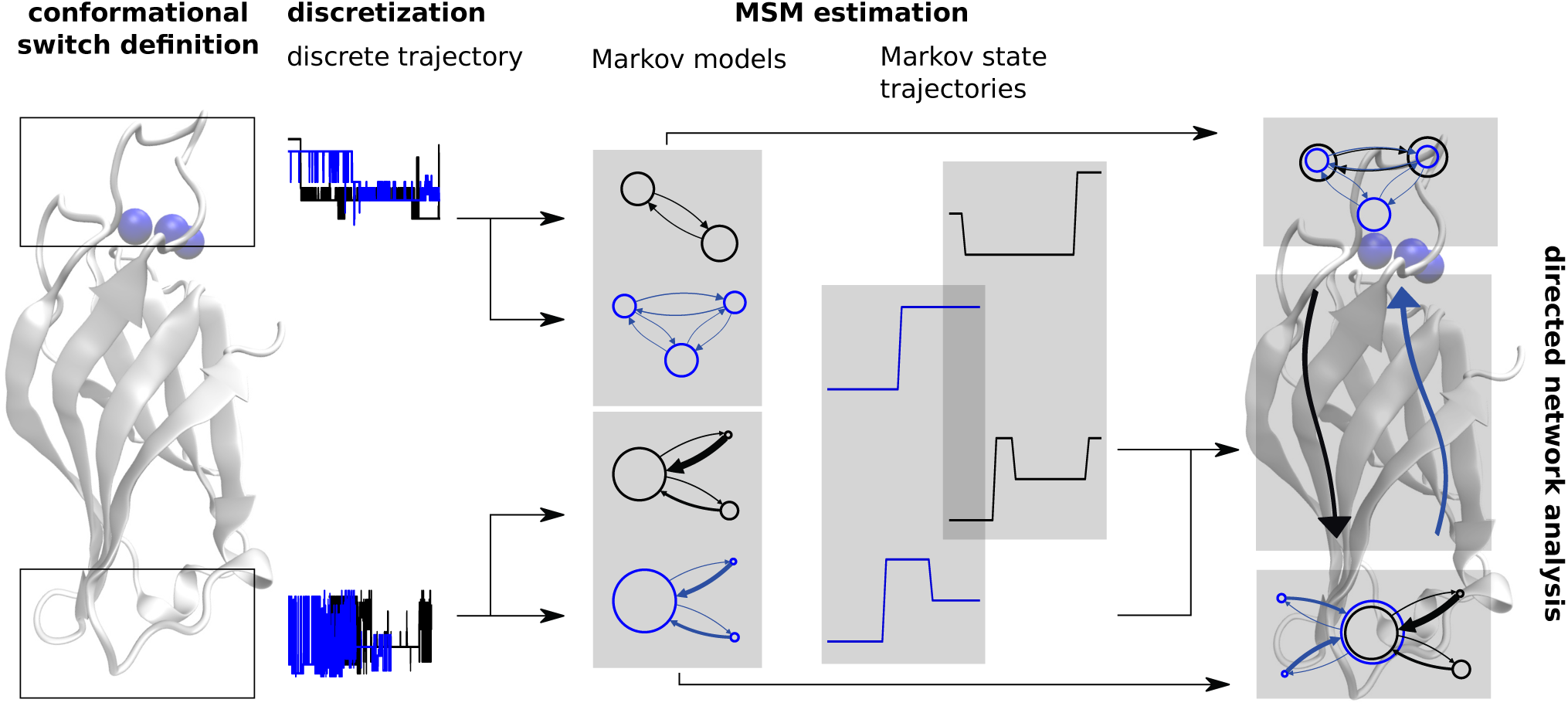
Method workflow at simplified example. From left to right: Two conformational switches are defined (top and bottom of protein) and raw MD trajectories of two ensembles (depicted by colors blue and black) are discretized into a (micro-) state space that is common to all ensembles of a particular conformational switch. Markov models are estimated separately, yielding transition matrices between metastable states and Markov state trajectories. In a following step, metastable states for each conformational switch can be identified between ensembles by comparing metastable state probability distributions, exploiting the common state space. Further, Markov state trajectories are used to estimate directional networks between conformational switches, exploiting the common time frame.

Our study is motivated by our interest for the role that protein conformational dynamics plays to determine spatio-temporal regulation of cellular processes and signaling. An abundant class of proteins that play important roles in cellular signaling (among other processes) is the C2 family consisting of 96,833 members^3^ of small beta-sandwich structured protein domains. C2 domains are membrane binders that in many cases require ion binding for activation.

One of the most exciting, yet unresolved problems connected to C2 domains is neurotransmitter exocytosis. Here, C2 domains play roles in vesicle recruitment (MUNC13) as well as in the actual release mechanism (Synaptotagmin-1, 2, 7 and 9).^4^ More specifically, synchronous neurotransmitter release is triggered by the double-C2-motif Synaptotagmin-1 (Syt-1) as a reaction to the electrically driven calcium concentration change. Syt-1 is specifically known to cause fast, concerted neurotransmitter release^5^ and hence is a good candidate for investigating calcium induced signaling with MD. Its structure and environment is summarized in Fig. 2. Even though the chain of reactions seems well understood, it is unclear how the ion concentration change triggers signaling on an atomistic scale. Possible mechanisms include a passive reaction due to the change in electrostatic potential, changes of protein structure or adapted conformational dynamics. While the first mechanism has been studied extensively, our aim is to investigate the role of Syt-1 conformational dynamics for its signaling function. Short preliminary test simulations of both Syt-1 domains, C2A and C2B, with and without membrane revealed that even single domain conformational dynamics is complex. Extensively sampling the whole system’s rare transition events between all possible membrane binding modes and protein conformations is prohibitively expensive with direct MD. We thus propose to break the problem down into obtaining kinetic models for individual components that will be coupled later.^1^ As a first step, we thus decided to analyze the Syt-1 C2A domain as it showed the most interesting behavior in our tests.

**Figure 2:**
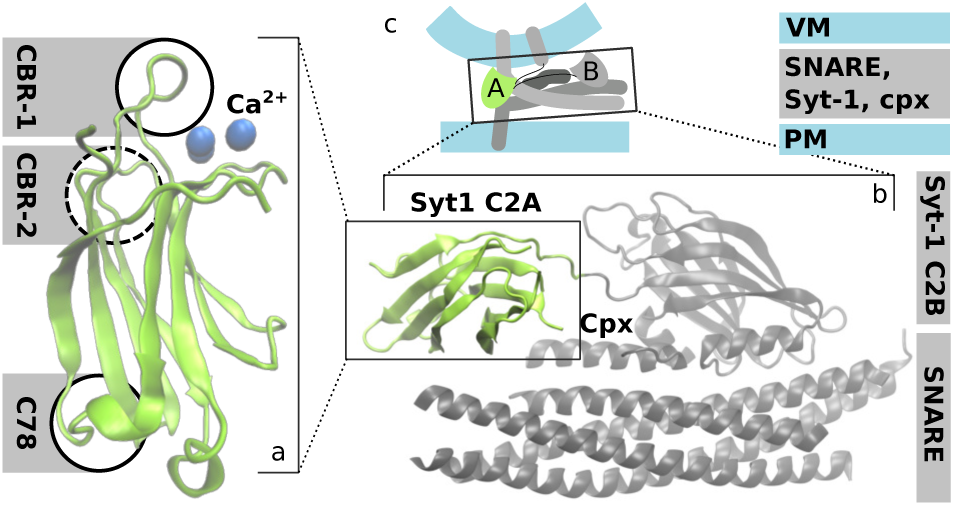
Syt-1 C2A with interaction partners for synaptic exocytosis. **a:** Structure of Syt-1 C2A, active sites used in later analysis highlighted by circles; calcium ions are depicted as blue spheres. **b:** Syt-1 C2A (green) in primed fusion complex with its sister domain, Syt-1 C2B, and SNARE/Cpx (gray) based on PDB 5W5C.^6^ **c:** cellular context with plasma membrane (PM) and vesicle membrane (VM). ^6^

The difficulties in understanding the inherently complex signaling process arise partially due to a lack of experimental methods capable of capturing the functional processes at a high spatial resolution over sufficiently long time. Molecular dynamics (MD) simulations are complementary here. They enable us to analyze C2 signaling function by assessing differences between the calcium bound and apo C2 domains using the Syt-1 C2A domain at atomistic resolution as an example. Markov modeling is then used to analyze the equilibrium reweighted conformational dynamics of the system, its Ca^2+^ configuration, and the calcium dependent information exchange between its different conformational switches.

## 2 Determining conformational switches and their coupling

We employ two main methods in order to analyze protein dynamics as a coupling of conformational switching domains: (1) Decomposing the protein structure into conformational switching domains and modeling their kinetics using local MSMs, and (2) Characterizing how these conformational switches are coupled by computing mutual information and transfer entropy between local MSMs. Here we outline the main concepts, see Fig. 1 for a visual summary and Methods for details.

### 2.1 Local Markov State Models identify conformational switches

Conformational switches are conceptually local protein regions that can exist in different discrete states, whose switching may be either spontaneous (i.e., by thermal noise), or induced by binding other changes of the equilibrium. Here we outline an approach to identify conformational switches that undergo spontaneous transitions in the simulation data.

We first decompose the protein or protein complex into conformational switching domains. When these are not obvious as in the present application, one can choose from a variety of methods to find optimal decompositions of the full system into sub-systems, e.g. Refs. ^7,8^

Secondly, we discretize the conformational switch state space into metastable states using MSMs.^9–15^ In the context of MD, MSMs model exchange between usually very fine-grained protein states by counting transitions between them. Based on this empirical count matrix, a transition probability matrix under reversibility constraints is estimated. Equilibrium probabilities of protein conformations follow from an eigendecomposition of this matrix. Not only does Markov modeling provide a means to combine arbitrary numbers of short off-equilibrium trajectories, but it also has turned out to be an efficient approach to learn about the slow and biologically relevant processes in vast MD data sets.^16–18^ We further note that kinetic modeling is necessary because in the finite data regime, MD datasets are no equilibrium samples. Thus, simple time averages in general do not represent thermodynamic ensemble averages. These can however be estimated using MSMs.

While various MSM approaches can be employed in order to identify metastable sets, such as Perron Cluster Analysis^19^ or VAMPnets,^20^ we here estimate local Markov models via the hidden Markov model (HMM) approach described in Ref.^21^ HMMs yield a mapping of the complex dynamics into an easily interpretable set of few “hidden” or metastable states of the local conformational switches. Besides the kinetic model, metastable trajectories (Viterbi paths, cf. Methods) are used for the analysis.

### 2.2 Coupling conformational switches via transfer entropy

To analyze pairwise directed influences between conformational switches, we exploit (a) the fact that metastable trajectories of different local protein features are time-synchronous, and (b) that the systems in the HMM formulation are Markovian by definition. We can thus simplify Schreiber’s original definition of transfer entropy^22^ using the HMM transition matrix *p*(*x*_*n*+1_|*x*_*n*_) and its stationary distribution *π*_*X*_(*x*_*n*_) that are defined for states *x*_*n*_ in a time series *X*. From time series *Y* to time series *X*, transfer entropy *T* (*Y* → *X*) hence reads

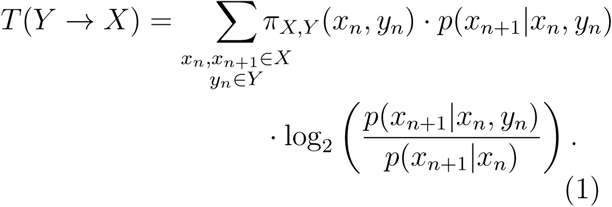

Joint probabilities in *X* and *Y*, *π*_*X,Y*_, are obtained from a transition matrix estimate in the combinatorial state space. The transition probability *p*(*x*_*n*+1_|*x*_*n*_, *y*_*n*_) is computed from the combinatorial transition matrix by marginalization over *y*_*n*+1_. This takes into account the common time frame and probes for causal effects of the current state onto the state that follows one Markov step in the future. Choosing the lag time of the underlying MSM ensures that our analysis indeed captures cross-talk of processes that our MSMs describe, even though other choices are theoretically possible.

Transfer entropy can be interpreted as a measure of the “incorrectness of [the] assumption” that “the state of [Y] has no influence on the transition probabilities on system [X]”. ^22^ It probes dependency in a direction-dependent fashion, i.e. is a heuristics for the amount of influence the process *Y* has onto the trajectory of a process *X*. According to this interpretation, we take *T* (*X* → *Y*) ≫ *T* (*Y* → *X*) as a statistical evidence for directed influence from process *X* to process *Y*.

## 3 Results

### 3.1 Syt-C2A consists of multiple metastable conformational switches

The analysis of conformational dynamics is conducted with focus on potentially functional protein regions. The Syt-C2A protein body adopts a *β*-sandwich fold which is naturally very rigid while the connections between the *β*-sheets are flexible. Furthermore, the largest flexible parts coincide with the calcium binding region (CBR), but there are several other flexible regions at various locations in the protein.

A simple analysis of root mean square fluctuations (SI) shows that only the CBR-1 loop becomes significantly stabilized with calcium binding. In the CBR-2, the opposite seems to happen, which hints towards an opening of the rigid body structure for this particular loop. CBR-2 is closely attached to the protein body in the crystal structure.

In order to obtain a human-readable model, we exploit our observation that Syt-C2A loops operate seemingly independent, and sub-divide Syt-C2A into three metastable regions of interest that are henceforth described as conformational switches: CBR-1, CBR-2 and a site opposite to the CBR (Fig. 2). The latter region will be called C78 since it connects *β*-sheets 7 and 8 (according to nomenclature of Ref.^23^). According to our analysis, the remaining regions express no metastability and will thus not be analyzed. We employ a Markov state model analysis, here using Hidden Markov Models (HMMs) in order to identify the metastable states within these conformational switches (Methods).

### 3.2 Local Markov models reveal calcium mediated loop dynamics

Our models predict that CBR-1 can form an *α*-helix as depicted in Fig. 3a. The free energy difference between the helical and other states is 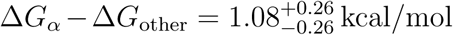 (without calcium) and slightly shifts to 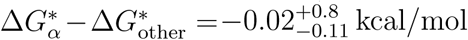 (with calcium). The life-time of this helical state becomes higher, from about 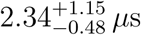 (without calcium) to 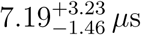 (with calcium). This means that the calcium binding enables a much more stable *α*-helical CBR-1 conformation.

**Figure 3:**
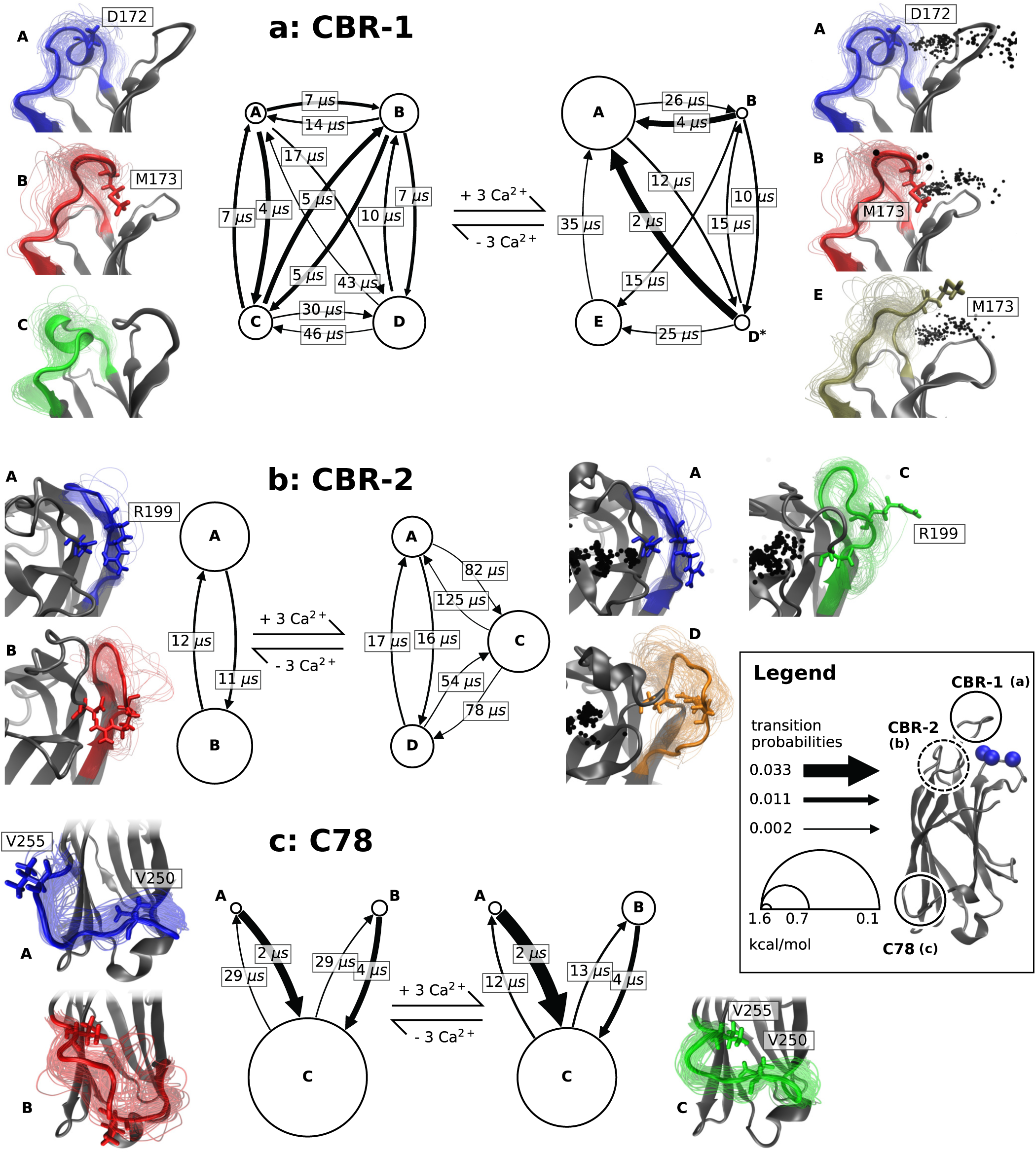
Markov models of Syt-1 C2A conformational switches in the apo state (left) and calcium-bound state (right). Capital letters denote metastable states. Representative structures are shown and color-coded to distinguish states within bound/apo models. Circle sizes are proportional to state’s equilibrium probabilities within each model. Arrows indicate transitions between metastable states, thickness proportional to the transition rate and annotated by pairwise inverse transition rate. **a:** MSMs of CBR-1 including sampled Ca^2+^ positions (black dots). States D and D* denote disordered structures that could not be assigned to unique structural elements. **b:** MSMs of CBR-2 with its loop in closed (crystal structure configuration, blue) and open (red, green, orange) conformations. **c:** MSMs of C78. Stick representations of single residues were drawn where instructive.

This population change is mediated by the three calcium ions inside the CBR which interact with the CBR-1 loops by Coulomb interactions, attracting the polar acidic residue D172 (Fig. 3a). This residue is located at the center of the *α*-helix sequence; Coulomb attraction towards the protein core stabilizes this structure. We corroborate this observation by noting that D172 has a very low solvent exposure in its *α*-helical conformation (SI). The second most probable structure in the calcium bound protein is a *β*-hairpin-like structure (Fig. 3a). In general, it is very rigid and gives high solvent exposure to M137.

However, even without calcium, stabilized configurations exist, as depicted in Fig. 3a). One of them is characterized by M173 being buried within the CBR (red structure in Fig. 3a). The stationary distribution seems tilted towards the M173-burying state in the calcium unbound protein. Besides a disordered state (denoted by D in Fig. 3a), CBR-1 can form another *α*-helix located towards the N-terminal end of this loop with D172 at its center (depicted in blue).

The CBR-2 loop in the crystal structures appears tightly bound to the protein body. Even though this conformation is populated in both our MD datasets, we find metastable states that are characterized by re-arrangement of CBR-2-stabilizing salt bridges (Fig. 3b).

In particular we note that the salt bridge R199-D178 which is found in the crystal structure and in all of the calcium-unbound trajectories (Fig. 3b, state A) can be opened by calcium binding. R199 becomes solvent exposed in this highest populated calcium bound state. This change goes along with a particular Ca^2+^ arrangement within the binding pocket (Fig. 4b,c). Mechanistically, the CBR-2 residues are influenced by the calcium ion charges. This explains why in the calcium bound trajectories, positively charged residues such as R199 can loosen from the protein body. The calcium binding thus enables R199 to be solvent accessible. Concerning the activation time, the conformational changes between the tightly bound state and the loosely bound ones is in the order of 12 *µ*s with one exception: In the calcium bound case, it takes 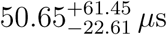 to attach the loop to the protein body again (result not shown in Fig. 3b).

**Figure 4:**
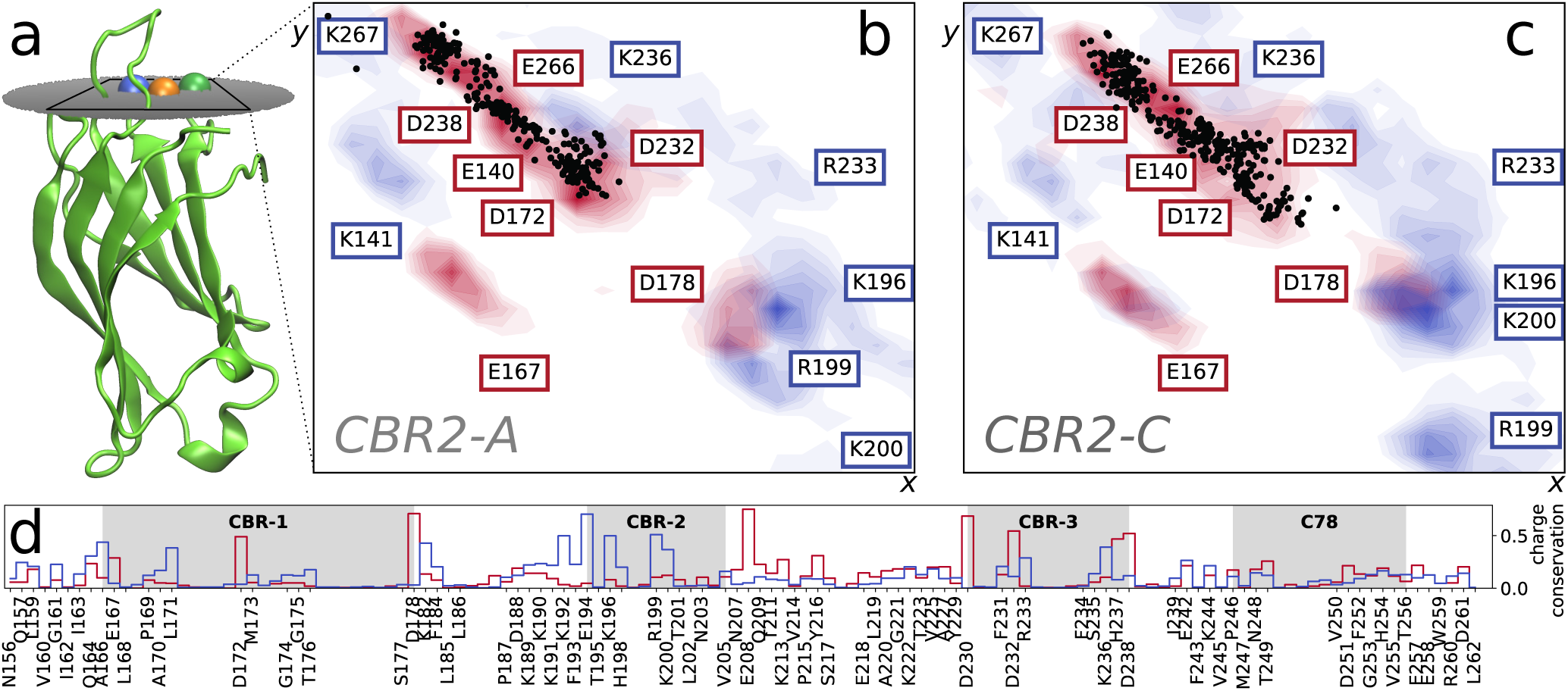
Ca^2+^ and binding pocket charge distribution. **a:** projection plane drawn in Ca^2+^-bound Syt-C2A structure. Panels **b** and **c:** Charge density averaged over 100 MSM samples and Ca^2+^ distribution in the calcium binding pocket. Ca^2+^ density is depicted by black points, positively charged amino acids as blue, negatively charged as red filled contour lines. MSM samples were drawn from CBR-2 MSM states A (panel b) and C (panel c), respectively. 2D projection (x, y) into the plane that is depicted in panel a. **d:** Conservation of charges among C2 domain family sequences (PFAM entry PF00168, seed alignment); histogram annotated with Syt-1 C2A domain residue sequence.

The C78 loop undergoes slow processes between three metastable states which could be identified in both, calcium bound and unbound data sets. With calcium binding, a shift of probability towards a V250-exposed state (cf. red structure in Fig. 3c) can be observed. Fig. 3c shows that two of the macrostates are characterized by releasing Valine residues from the protein body. These are V250 and V255 which can be attached independently of calcium binding. In order to switch between the V250 to a V255 exposing state, both residues must be attached to the protein body as an intermediate state.

### 3.3 Ca^2+^ distribution depends upon protein conformation

When analyzing the calcium configuration within the binding pocket we find that the Ca^2+^ distribution relaxes from the crystal structure positions into a broader distribution due to Coulomb repulsion between the ions. The crystal structure position of Ca1 acts as the inner edge of the binding pocket which is structured like a funnel along which the ions can collectively move. Therefore, Ca^2+^ distribution is dynamic and is concentrated across multiple possible binding sites that are more or less defined. The residues forming these binding sites are dynamic themselves, i.e. take different conformations in different metastable states as exemplified in Fig. 4b,c.

The importance of the charge distribution created by binding pocket residues becomes evident with a sequence alignment of all known members of the C2 family.^3^ The histogram of conserved charges derived from the sequence alignment (Fig. 4 d) confirms that charges at Ca^2+^ coordinating residues (Syt-1 C2A residues D172, D178, D232, D230, D238, also reported by Ref.^24^) are indeed conserved. Further, we find that positive charges at position 199-200 (Syt-1 nomenclature) are conserved, underlining the significance of our findings. Another distinct feature in the charge conservation histogram is a conserved region of positive residues around K190, which corresponds to the Lysine rich cluster described by Ref. ^25^ for the C2 domains of Syt-1 and rabphilin 3A.

We can further qualitatively generalize our results. Beyond the sequence conservation, C2 domains are highly conserved in structure as the resolved structures from the PFAM C2 family^3^ have an RMSD of 1.80 ± 0.36Å to the analyzed crystal structure (1BYN). ^26^ It thus becomes very likely that conformational dynamics plays a role for other C2 domains, too, most likely connected to CBR (or equivalent for non-calcium binders) or other non-*β*-sheet structural elements such as C78.

### 3.4 Distant protein features show allosteric behavior

In Markov state modeling and other statistical analysis methods, there is a trade-off between the complexity of the system to analyze and the amount of detail one can resolve with statistical significance. The presented Syt-C2A example has distinct regions in which conformational switches occur. As commonly observed, obtaining a joint MSM including all combinations of conformational switches proved difficult due to insufficient statistics.^1^ However, high-quality MSMs could be obtained for individual conformational switches, in the field of the remaining protein. Here we investigate how these different sub-systems interact using an information theoretic approach to model the network.

A qualitative assessment of un-directional influences using Pearson’s correlation coefficient (of coordinates) and mutual information (of MSMs) shows that there indeed is a weak coupling between conformational switches. As expected, coupling between spatially adjacent sites is higher than between distant sites. In particular, Ca^2+^ ions tighten the coupling between CBR-1 and CBR-2 (Fig. 5).

**Figure 5:**
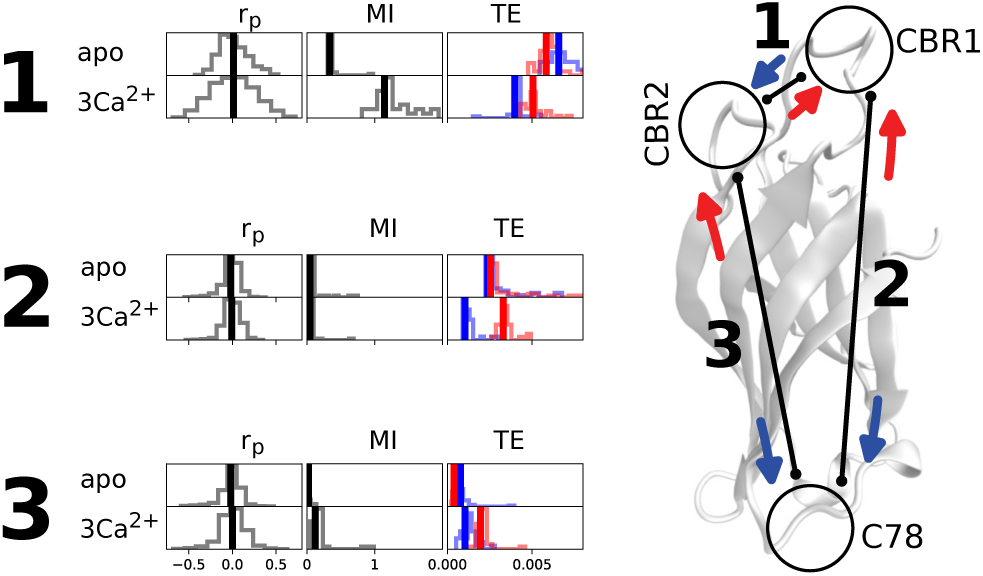
Coupling between conformational switching domains. Each link between conformational switches (enumerated 1-3) is quantified by naive computation of Pearson’s correlation coefficient between plain coordinates (*r*_*P*_), mutual information (MI) and probed for directionality with transfer entropy (TE). TE color code refers to different directions as shown in crystal structure. Bootstrapping histograms for MI and TE are shown to assess the estimation error (MI, TE columns). The correlation coefficients between all pairs of heavy atoms of two protein regions are depicted by histograms to qualitatively assess linear couplings (*r*_*P*_ column).

It is only natural to ask if this coupling is symmetric or if any kind of uni-directional influence between conformational switches can be inferred. A crucial point for the analysis is that a common time frame can be defined from the Markov state trajectories of the individual conformational switches as they occur simultaneously. Transfer Entropy *T* (*X* → *Y*) is a well established measure for assessing directional networks between time series *X* and *Y*.^22^ It can be interpreted as a statistical measure for the amount of information transported between two systems and, thus, is direction dependent. We say that *X* influences *Y* if *T* (*X* → *Y*) ≫ *T* (*Y* → *X*) (Methods). This attempts to model statistical *causality* (in the Granger-sense) between local protein features. In Granger’s original definition,^27^ in simple words, a random variable *X* “causes” *Y* if knowledge of the past of *X* reduces the uncertainty of predicting *Y* ‘s future. Transfer entropy relates to this concept; ^28^ therefore we apply a tool corresponding to a very broad definition of allostery.

The transfer entropy estimates from our data are presented in Fig. 5. We can identify a weak non-symmetric coupling between C78 and CBR-1 in the calcium bound case. The transfer entropy between this pair of loops is significantly higher from C78 to CBR-1 than vice versa, i.e. the prediction error for CBR-1 dynamics can be decreased by adding information about C78. This can be interpreted as a causal influence from C78 onto CBR-1, suggesting an allosteric coupling that is induced by calcium binding. It goes along with a slight increase in mutual information between this pair of conformational switches. We hypothesize that the effect is caused by long-range Coulomb interactions from the ions that is further mediated by the network of chemical bonds of the rigid protein body.

Please note that this result is purely based on statistical analysis and, however desirable, no mechanistic interpretation follows here.

## 4 Discussion

We have shown that challenges arising from the MD sampling problem and the curse of dimensionality can be met by combining local Markov state models of conformational switches with an information-theoretic analysis of their coupling. We find that the estimation of few-state local Markov models is very robust and avoids commonly experienced statistical problems with estimating Markov models of the global protein state. Further, local models are easy to interpret. We therefore think that the present approach will be instrumental for the modeling of large proteins with loosely coupled local conformational switching domains.

Our results demonstrate that the effect of calcium binding to Syt-C2A is not a simple linear response as might be expected by increasing the charge in a specific part of the system, but rather a complex change of the system kinetics. Even though experimental^29^ and MD studies^30,31^ suggested that Syt-1 C2A does not switch between well-defined conformations upon calcium binding, sampling the system extensively shows that the C2A domain indeed undergoes subtle and very rare conformational switching which is profoundly impacted by Ca^2+^.

Further, it is well established that upon calcium binding, the C2A domain effectively switches off its membrane repelling charge,^32,33^ thus being attracted by the membrane while interacting with the SNARE-cpx complex (cf. Fig. 2 and Refs.^5,6^). This model however does neither specify such an interaction nor does it explain how the signal (calcium ions) spreads to possibly distant interaction sites. Our model suggests that the conformational dynamics is altered upon calcium binding. This could be a potential mechanism for the calcium signal spreading to distant sites.

As any study based on MD simulations, this work relies on the forcefield that was used to generate the data. In particular, we note that the applied calcium ion model is approximative and e.g. does not account for electronic screening of charged moieties or environment-dependent changes in partial charges.^34,35^ Even though this should be taken into account when basing experiments on our results, we are confident that charge-mediated effects such as the ones laid out in this work can be described nevertheless. Comparison to other forcefields, in particular polarizable ones, would improve the prediction quality but is beyond the scope of this conceptual work.

At this point, our system does not include elements of the cellular environment such as phospholipid membranes. We note, however, that the presented methodological framework would be applicable and could be used to quantify effects of membrane binding to C2A conformational dynamics.

Our model predicts a calcium induced population change towards a CBR-1 *α*-helix (Fig. 3a), burying negatively charged residues such as D172, thereby making the CBR more attractive to the membrane. This is consistent with the charge neutralization argument of Ref.^32^ Membrane attraction is further enhanced by hydrophobic residues that are not necessary for stabilizing the *α*-helix. The CBR-1 configurations that, according to our model, are highly populated in the calcium unbound protein (Fig. 3a) might even reinforce the membrane repelling function by moving hydrophobic residue M173, which is exposed in the disordered CBR-1 configuration, into the binding pocket. CBR related polar acidic residues are solvent accessible in this case since they are not bound by calcium ions. This has the side effect that CBR attraction to solvent calcium ions rises.

In the CBR-2, we find that calcium binding enables release of R199 from the protein body (Fig. 3b). As R199 has been reported to be of importance for membrane penetration,^23^ ionic interactions with SNARE,^36^ and forming the C2A-C2B interface,^37^ our model might be key for understanding the interactions with the fusion machinery. In particular, Syt-1 interaction with SNARE might be triggered by calcium that, given its positive charge, induces the release of R199.

The C78 region generally contains many hydrophobic amino acids which opens the possibility of potential weak membrane interaction at this non-CBR site, a model previously proposed for Syt-1 C2B.^38^ This interaction would allow the C2A domain to bind both, vesicle and plasma membrane, at the same time. Unfortunately, we are not aware of thorough studies that investigate C78 or adjacent sites. In order to verify our model, we thus propose experimental validation. Such a validation would include assessing the relevance of this protein site e.g. by knock-out mutations of all hydrophobic residues located there. If effects on membrane fusion can be determined, singlet mutants could be used to validate our predictions. We expect especially Valine residues 250 and 255 to alter functionality since their configurations are characteristic for metastable states in the local models.

Our study further suggests cooperativity between the loops. As expected, our correlation and mutual information analysis shows that CBR-1 and CBR-2 loops are weakly coupled. Our model further predicts that calcium binding induces a stronger coupling. Transfer entropy analysis suggests that calcium binding increases the influence that C78 exerts onto CBR-1, even though the absolute magnitude might be rather low. This weak allosteric coupling insinuates that perturbation of C78, e.g. through membrane contact or binding to another protein in the fusion complex, effects on the conformational dynamics of CBR loops. Even though our model does not predict the properties of such an allosterically induced change, we believe that this effect should be taken into account in further studies of C2 domains.

Transfer entropy analysis as conducted in this work is limited to model the interplay of conformational switches. Modeling a high resolution allosteric pathway with this technique is however possible and subject of current research.

We can further qualitatively generalize the mechanisms described here for Syt-1 C2A to a larger spectrum of proteins that are not necessarily connected to neurotransmitter secretion. As sequence alignment of the charges in the C2 family (Fig. 4 d) has shown, especially our newly described calcium dependent mechanism of CBR-2 regulation might indeed be a structural feature of C2 domains in general.

The present approach relied on identifying the conformational switching domains treated by local Markov models with expert knowledge. The development of machine learning approaches to select these conformational switches automatically is a topic of future research.

## 5 Methods

### 5.1 Molecular dynamics simulations

In order to observe all important dynamical processes of the system, we have carried out extensive sampling. 220 trajectories of 2 *µ*s individual length have been generated, adding up to a cumulated simulation time of 440 *µ*s, which to the best of our knowledge is the largest atomistic MD dataset generated for a C2 domain up to now. Since synaptic vesicle fusion can be triggered by calcium in less than 100 *µ*s^4^ the assumption that all relevant processes have been sampled is justified.

Simulations have been carried out for the Syt-C2A domain in its Ca^2+^-bound and Ca^2+^-free state using the CHARMM36 force field.^39^ The setup is based on the C2A structure contained in PDB 2R83 (which contains the double motif C2AB) and Ca^2+^ ion positions are initiated from PDB 1BYN which contains three calcium ions bound to the CBR. Starting structures for MD production runs were randomly drawn from a smaller precursive dataset. For both setups a water box of side length 6 nm was generated with a KCl ion concentration of 0.1 mol*/*l at neutral total charge using the TIP3P^40^ water model. The setups contain 8363 (calcium free) and 8359 water molecules (with calcium), respectively. The setup as well as equilibration in the NVT and NPT ensembles (100 ps each) were conducted with Gromacs,^41^ production run simulations of 2 *µ*s were carried out in the NPT ensemble at 300K and 1 bar using the OpenMM software package.^42^ We used 1 nm cutoff for non-bonded interactions, PME electrostatics, and rigid water molecules. We further exploited the heavy hydrogen approximation as described in the openMM documentation, allowing a 5 fs integration step with the openMM Langevin integrator. An aggregated simulation time of 184 *µ*s was obtained for the calcium free case, 256 *µ*s for the calcium bound case, in each case the accumulation of new data was continued until converged MSMs were obtained. Visual representation of molecular structures were obtained with VMD.^43^

### 5.2 Local MSMs

For modeling local loop dynamics, the full (bound and apo) trajectory data was directly clustered according to a minRMSD norm. This includes superposing the full protein (using MDTraj^44^) and defining discrete states according to the Euclidean norm between all atom coordinates of a specific region. This includes amino acid side chains and hydrogen atoms. We chose A170-T176 (CBR-1), R199-N203 (CBR-2) and T249-E258 (C78) to extract the local conformations, respectively. 15-30 discrete states and the *k*-means algorithm^45^ were used for each local model. The selection of regions is inspired by a precursive full protein analysis using pairwise minimal residue distances between blocks of two residues and a state discretization in a low dimensional TICA projection.^46^ Besides a relatively low spatial resolution at local sites, this analysis did not yield a valid MSM due to disconnected combinatorial states. It however motivated us to conduct analyses on local sites and to select the ones with metastable dynamics.

Discretizing local protein features after global minRMSD superposition could potentially yield spurious correlations between distant protein sites.^47^ We note that this is unlikely because a) the largest mass of the protein is concentrated in its rigid *β*-sandwich body and b) the comparably low number of discrete states at local sites is too small to capture such subtle influences. Microstate definitions at example of CBR-2 are shown in the SI. Further and most importantly, the hidden metastable states that we discuss and use for mutual information and transfer entropy analysis reflect major internal loop rearrangements and are therefore most likely not affected by superposition artifacts.

HMMs were estimated separately for calcium bound and unbound data sets based on the common discretization described above. In contrast to Markov state models, HMMs are not based on the assumption that the microstates obtained by clustering are approximately Markovian. Instead, a number of hidden states (usually less than the observed microstates) are introduced as explained in detail in Ref.^48^ The number of hidden states was chosen such that it yields the best resolution of the slow processes in the protein and fulfills the validation criteria for Markov modeling (see below). All HMMs were estimated at lag time 50 ns using the PyEMMA software package.^45^ We observed that only in relatively small configuration spaces (such as the ones defined by partitioning the protein into conformational switches), a robust, common discretization into Markov states could be obtained.

Corresponding HMM states between calcium bound and unbound HMMs were identified by computing KL-divergences between their observation probabilities per microstate (i.e. in the joint state space) and using a cutoff to identify hidden states across datasets (cf. SI). This operation is only possible because all data is discretized the same way i.e. microstate definitions do not differ. We use the KL-divergence between these distributions, an example (identification of macrostates for C78) is shown in the SI. Note that the KL-divergence becomes zero in the limit of equal distributions, thus one can define a cutoff of 2 below which states are identified for all models.

As our HMMs display high metastability, we can use the approximation *K* = *T* − 𝟙 to compute the rate matrix *K* from the transition matrix *T*. The elements of *K* are pairwise inverse transition times.

Markov models were validated using two measures.^15^ First, the implied time scales were tested for convergence. We further tested for stationary distribution convergence to ensure that the presented results do not depend on the model lag time. Second, the Chapman-Kolmogorow equation was used to check for consistency between predictions of the presented models with direct estimates at higher lag times. Using Bayesian sampling of the posterior, the error was estimated for all the quantities computed by the MSMs/HMMs.^49^ Results are presented in the SI.

### 5.3 Free energy validation of ion forcefield

For model validation purposes, we derive binding free energies of each Ca^2+^ using alchemical free energy perturbation methods^50,51^ for the crystal structure conformation.

As a buffer solution contains monovalent ions, it can be expected that calcium binding sites are normally saturated by those ions. This is confirmed by our calcium-free simulations in which we observe Potassium ions in the binding pocket. Alchemical free energy perturbation was hence simplified to a computational transformation of Ca^2+^ → K ^+^ in the binding pocket and the opposite transformation in the solvent.

We further assume that in experiments, ions fill the binding pocket subsequently, i.e. we compute Ca3 in the presence of (Ca1, Ca2), Ca2 in the presence of Ca1, and Ca1 by itself. As we are interested in the binding free energy difference of a particular protein conformation, we apply harmonic constraints to the full protein. Relative free energies were computed using MBAR.^51^

Our results were validated according to Ref. ^52^ This includes the validation of alchemical intermediate overlap and convergence analysis. Further, we repeat the same calculation at least 20 times to make sure that our results are reproducible.

The resulting ion binding free energies from the equilibrated crystal structure are comparable to experiment within error, however do not fully match the experimentally observed order (SI).

### 5.4 Combinatorial Viterbi path model

Since a central concept of this work are hidden state space trajectories or Viterbi paths, the most important concepts^48^ are summarized here. The Viterbi algorithm maximizes the probability *P* of the hidden state sequence *q*_1_*q*_2_ … *q*_*n*_ with the observed sequence *O*_1_*O*_2_ … *O*_*n*_ given our model *λ*

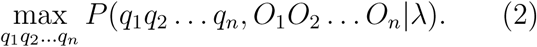

To this end, the Viterbi path is a maximum likelihood path on the hidden states. It naturally comes with the time step of the underlying HMM. Its estimation algorithm is also part of PyEMMA.^45^

The degree up to which the local models communicate was estimated by comparing the stationary distribution of independent models to a new estimate. By assuming independence, this property follows directly from the product of local model probabilities per state.

Without assuming independence, a new estimate for the dynamics in combinatorial state space was made based on the local model Viterbi paths. It yields the combinatorial transition matrix or conditional probabilities *p*(*x*_*i*+1_, *y*_*i*+1_|*x*_*i*_, *y*_*i*_) and joint stationary distribution *π*_*X,Y*_ (*x, y*) of two processes *X* and *Y*.

### 5.5 Mutual information

Let *X* and *Y* be Markov processes that are localized at distinct spatial features of a protein and are well-described by (local, individual) HMMs. The mutual information *M* of *X* and *Y* is defined in terms of the joint probability *p*_*X,Y*_ (*x, y*) and the independent probabilities *p*_*X*_(*x*), *p*_*Y*_ (*y*) of *x* ∈ *X* and *y* ∈ *Y*. In this work, we identify these probabilities *p* as the stationary probabilities of Markov models, most commonly denoted by *π*. We can thus write

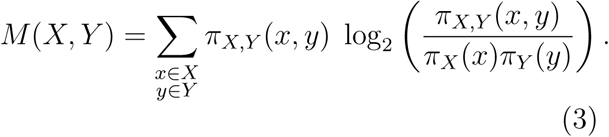

We obtain the independent stationary distributions *π*_*X*_, *π*_*Y*_ from the local hidden Markov models and the joint stationary distribution *π*_*X,Y*_ from a combined Markov model estimated based on the Viterbi paths. We use Eq. 3 to compute mutual information between all pairs of conformational switches of Syt-1 C2A.

For interpreting mutual information, it can be re-written in terms of the KL-divergence **D**(·‖·) as follows:

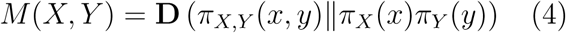

Mutual information can thus be interpreted as a measure of the incorrectness of the assumption that both systems are independent.^22^

## Acknowledgement

Funding from DFG SFB / TRR186 (project A12) is gratefully acknowledged. We further thank the Einstein Foundation Berlin (project SoOPic to N.P.) and the European Commission (ERC CoG 772230 “ScaleCell” to F.N.). We are grateful for discussions and feedback to Martin Scherer, Simon Bärfuss, Thomas Söllner, Fabian Paul, Shreyas Kaptan, Sebastian Stolzenberg, Guillermo Pérez-Hernández, Katarzyna Ziółkowska, Andreas Mardt.

## Supporting Information Available

Method validation measures as well as further information about the Syt-1 C2A system are provided in the SI.

