## Supplemental Material for "Coupling of conformational switches in calcium sensor unraveled with local Markov models and transfer entropy"

#### 1 Additional Syt-1 C2A analyses

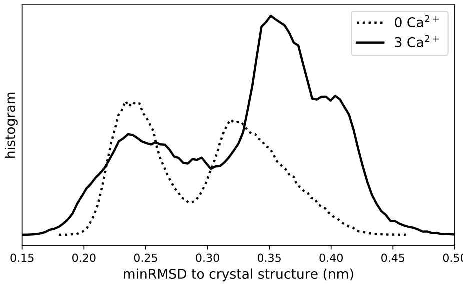


**Figure 1:** Histograms of minimum root-mean-square deviation (RMSD) between calcium bound and unbound trajectories to their respective PDB structures (1BYN: bound; 2R83: unbound). The multi-peak structure shows that besides the crystal structure, reasonably populated conformations exist which become more populated in the bound case.

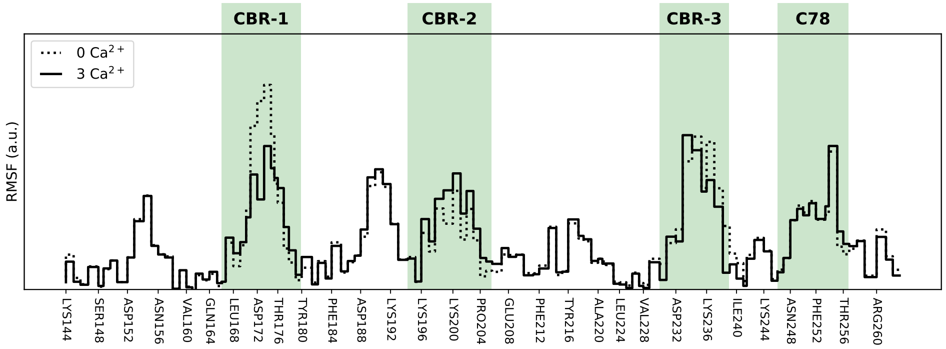


**Figure 2:** Syt-C2A RMSF per residue as computed from the full set of trajectories. Several regions appear to have significant motion, most of them correspond to CBR loops.

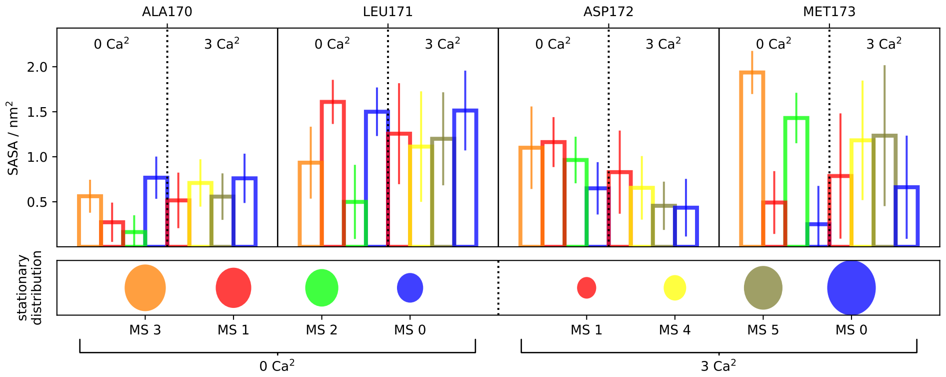


**Figure 3:** SASA of charged polar or hydrophobic residues in CBR-1 (upper panel). The bars are color-coded according to the convention used throughout this paper. Stationary probabilities are added (lower panel). SASA and stationary distributions of calcium bound and unbound states are sorted left and right of the vertical dotted lines for each residue.

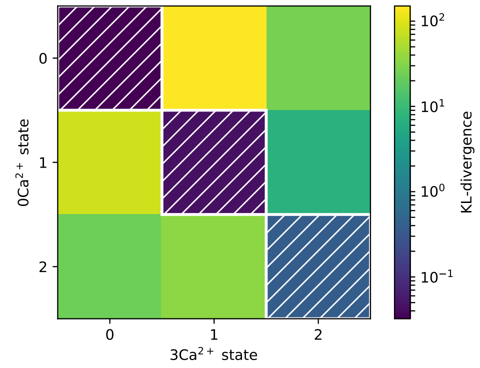


**Figure 4:** HMM macrostate identification: KL-divergences between state observation probabilities per macrostate for calcium bound and unbound datasets. Both HMMs have 3 hidden states. White hatches depict states that are identified between datasets.

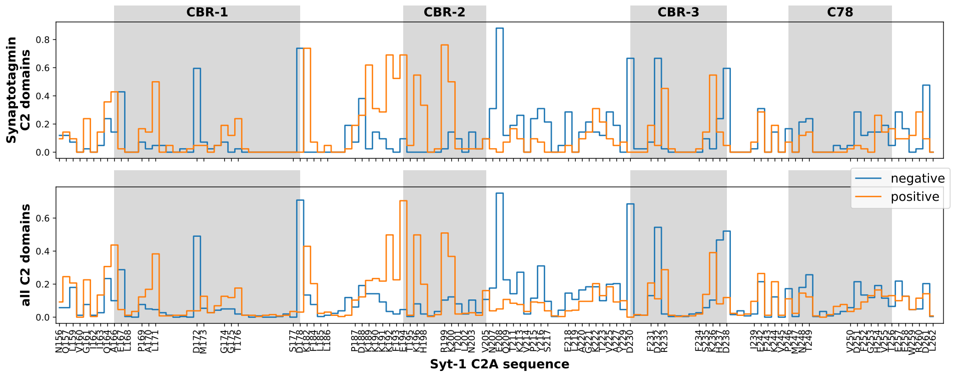


**Figure 5:** Bottom: Sequence alignment of C2 domain family members (PFAM entry PF00168, seed). Bottom: Sub-sample of synaptotagmin C2 domains.

### 2 Method supplement

#### 2.1 Local clustering

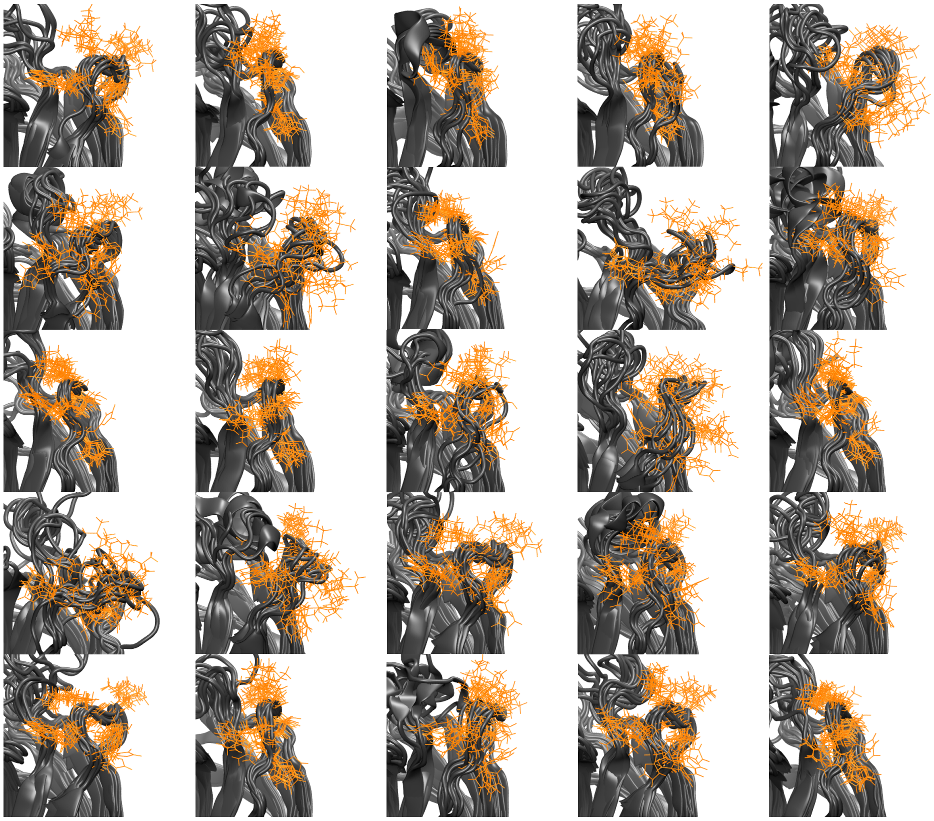

**Figure 6:** Representative structures of microstates of CBR-2 obtained from k-means clustering. Residues AR199-N203 that were used for discretization are color coded in orange. Microstates mirror internal configurations of CBR-2 as well as its distance to the protein body.

#### 2.2 Markov model validation

Generally, when building a Markov model, it needs to be checked if the process in discrete state space is Markovian for a certain lag time $\tau$. This can be done by checking for the convergence of model properties with respect to the model parameter $\tau$ [2] and by testing for the Chapman-Kolmogorov equation.

**Implied timescales and stationary properties convergence.** When sampling rare processes in a large data set however, one faces the problem that processes can be lost at high time scales. This is why the model parameter was tuned to a value which allows the implied timescales and the stationary distribution of all six models to be constant within error.

As depicted in Fig. 7, especially in the case of calcium bound CBR-1, the interval between timescale convergence and loosing the process is very short. The reason is that this rare event is sampled poorly. Nevertheless it is assumed that this model is valid since the chosen lag time of $\tau = 50\,\mathrm{ns}$ ensures Markovianity in all of the other models. Further, the behavior is reflected in the error estimate of the derived properties such as stationary distribution and mean first passage times (MFPT).

**Chapman-Kolmogorov test** Consistency of the Chapman-Kolmogorov equation $T(n \cdot \tau) = T(\tau)^n$ is tested for all the models presented here. As already mentioned, we are operating in the data sparse regime, so estimates can only be done for a finite number of multiples of the lag time $\tau$.

Predictions as well es estimates from the presented models are depicted in Fig. 8. Generally, we note that transition probabilities have the same trends, i.e. stay in the same order of magnitude. The presented error is estimated by Bayesian sampling of the posterior, it shows significant overlap in all cases. However, data sparsity leads to divergence at lag times of $3 \cdot \tau = 150\,\mathrm{ns}$.

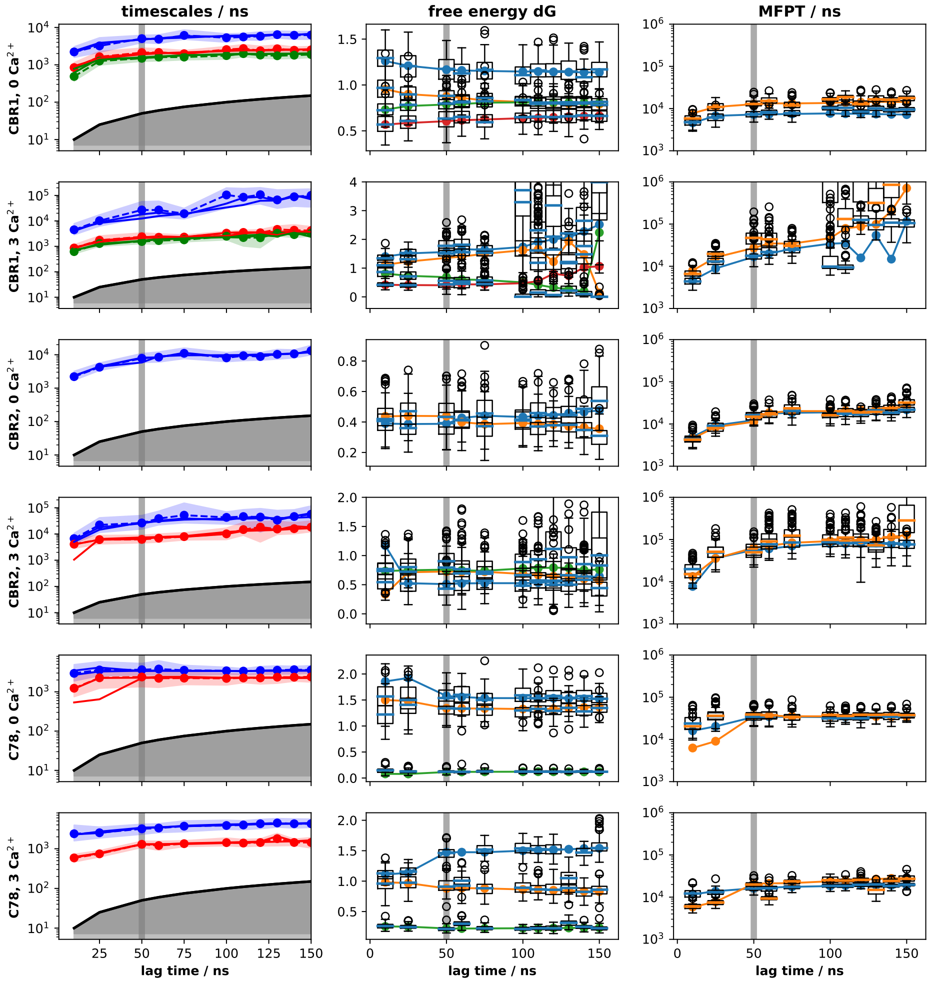


**Figure 7:** Convergence of implied timescales (left column), relative free energy (middle) and mean first passage times between arbitrarily chosen states (right).

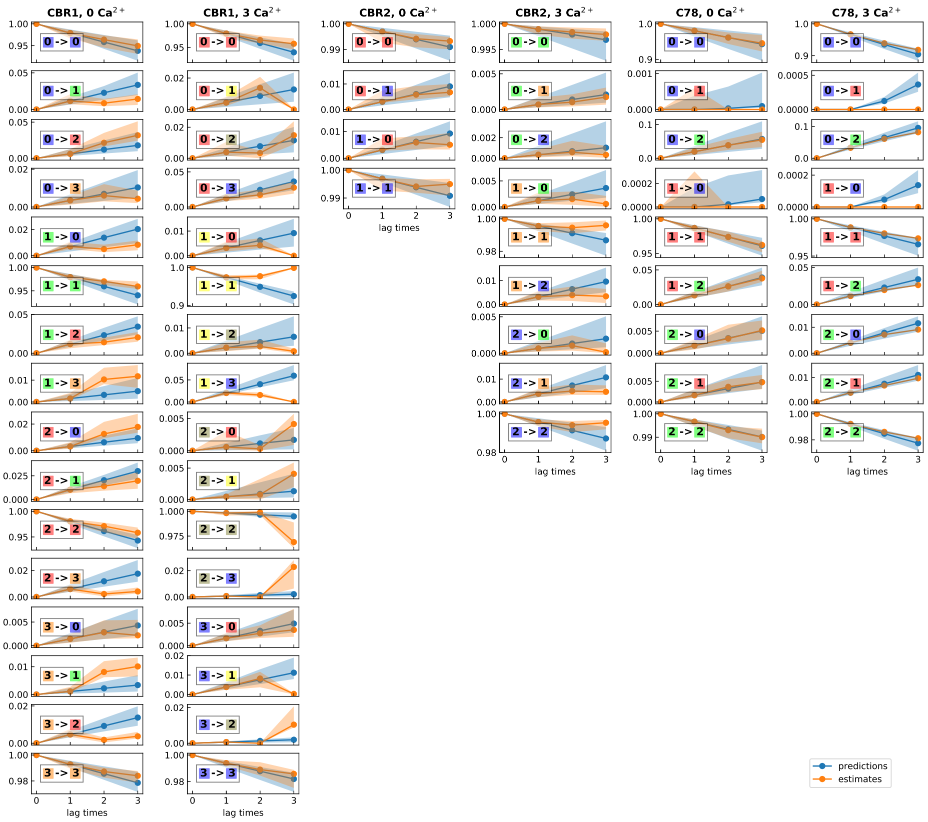


**Figure 8:** Result of the Chapman-Kolmogorov test column-wise ordered by model. Predictions from the presented models are depicted in blue, new estimates in orange. Transitions are encoded in macro state numbers and colored as in the results section.

#### 2.3 Validation of transfer entropy and mutual information

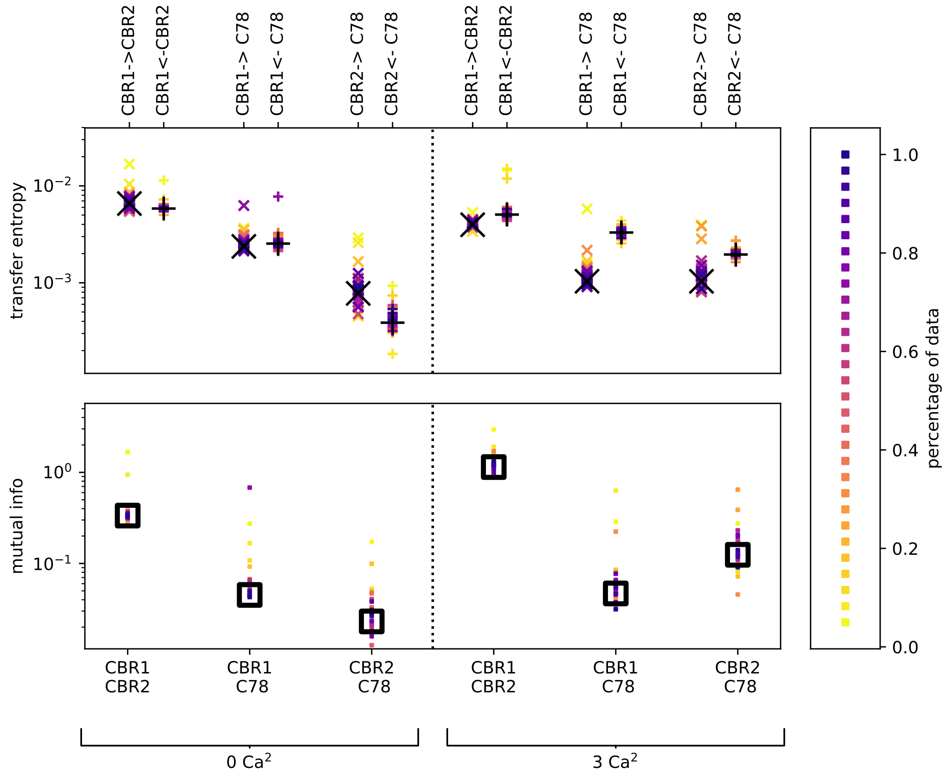


**Figure 9:** Bootstrapping validation for mutual information (bottom) and transfer entropy (top). Estimates using the full set of data are depicted with large black symbols. Left panels show calcium unbound, right panels calcium bound data.

Mutual information and transfer entropy were validated using bootstrapping of trajectory data (cf. Fig. 9). In order to assess if the results are significantly different to zero, a comparison was made to shuffled trajectories, i.e. the time information within the trajectories was kept constant while trajectories were combined that did not happen at the same time. As depicted in Fig. 10, the results in this case are at least one order of magnitude smaller than the results in the correct time frame.

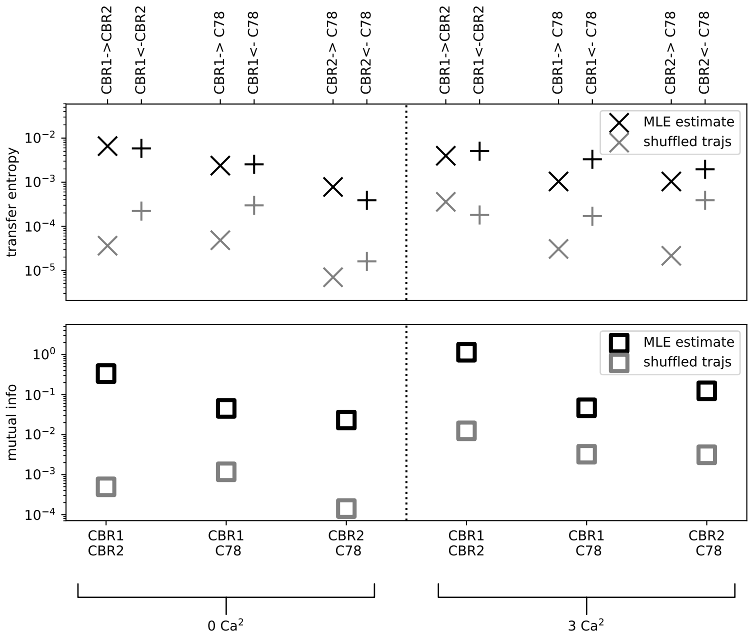


**Figure 10:** Mutual information and transfer entropy validation by comparing to results obtained from shuffling trajectories among each other but keeping frames within the single sub-system trajectories.

#### 2.4 Ion model validation by Alchemical free energy perturbation

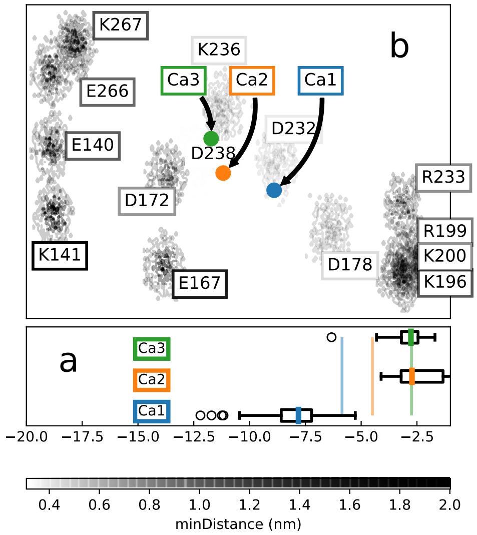


**Figure 11:** Alchemical free energy computations and binding pocket projection for crystal structure. Box plots of free energy results shows that experimental values (solid vertical lines) can be matched within the error. The only exception is ion Ca2 which in terms of its binding free energy is indistinguishable from Ca1. Configuration of (color coded) calcium ions is shown within binding pocket of crystal structure.

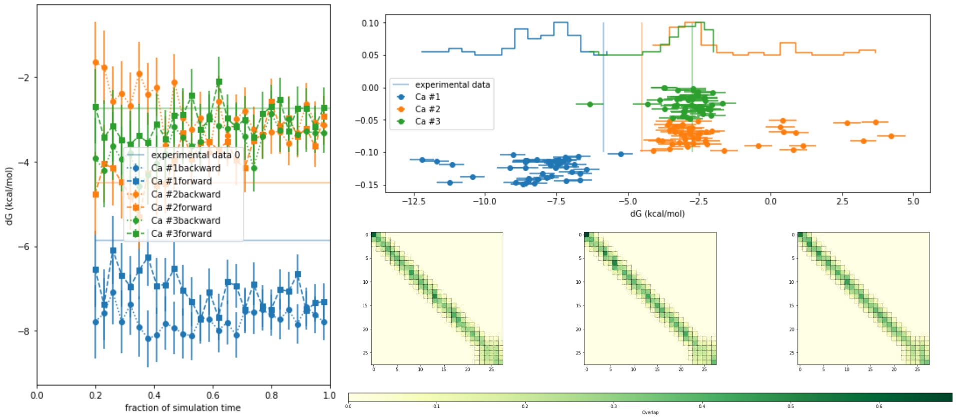


**Figure 12:** Validation of alchemical free energy computations with MBAR using crystal structure. Left: Forward-backward convergence. Top right: results of about 50 indipendent computations. Bottom right: Overlap matrices with boxes denoting sufficient overlap according to Ref. [1].
